# Nanoparticle-Functionalized Acrylic Bone Cement for Local Therapeutic Delivery to Spinal Metastases

**DOI:** 10.1101/2023.02.06.527220

**Authors:** Ateeque Siddique, Megan E. Cooke, Michael H. Weber, Derek H. Rosenzweig

**Affiliations:** Department of Surgery, Division of Orthopedic Surgery, McGill University and the Research Institute of the McGill University Health Centre, Injury Repair & Recovery program, Montreal, QC, Canada

**Keywords:** PMMA, mesoporous silica nanoparticles, chemotherapeutics, local delivery, 3D culture

## Abstract

Polymethylmethacrylate bone cement is often used to reconstruct critical-sized defects generated by surgical resection of spinal metastases. Residual tumor cells after a resection can drive recurrence and destabilization. Doxorubicin (DOX) is a common chemotherapeutic drug with unwanted side-effects when administered systemically. Mesoporous silica nanoparticles are gaining attention for targeted drug delivery to bypass the negative side effects associated with systemic drug administration. We developed a nanoparticle-functionalized cement for the local release of DOX and tested its ability to suppress cancer cells. DOX was loaded onto nanoparticles which were then mixed into the cement. Drug release profiles were obtained over a period of 4 weeks. Cement constructs were incubated with 2D and 3D cultures of breast and prostate cancer cell lines, and cell metabolic activity and viability were evaluated. Cell migration and spheroid growth were assessed in collagen-coated spheroid cultures. Nanoparticles were homogenously dispersed and did not alter cement mechanical strength. A sustained DOX release profile was achieved with the addition of nanoparticles to the bone cement. The release profile of DOX from nanoparticle cement may be modified by varying the amount of the drug loaded onto the nanoparticles and the proportion of nanoparticles in the cement. Cells treated with the cement constructs showed a dose- and time-dependent inhibition. Cell migration and spheroid growth were impaired in 3D culture. We show that nanoparticles are essential for sustained DOX release from bone cement. DOX-loaded nanoparticle cement can inhibit cancer cells and impair their migration, with strong potential for *in vivo* translation studies.

## Introduction

The spinal column is the most common skeletal location of cancer metastasis. Spinal metastases occur in 20-40% of all cancer patients, originating primarily from breast, prostate, lung, and kidney cancers [1–3]. About 20% of spinal metastases patients are symptomatic, including pain and neurological dysfunction due to instability and epidural spinal cord compression (ESCC). Spinal metastases result in lesions in the affected vertebrae which can cause pathologic compression fractures due to the decreased load-bearing capacity of the spine [1, 2, 4]. The prevalence of spinal metastases is expected to increase with patients living longer due to improving detection and medical/surgical treatment for many cancers [4]. Furthermore, the overall postoperative survival for patients undergoing surgery for spinal metastases has improved significantly in the last 20 years [5]. Thus, it represents an increasingly important oncologic challenge. The treatment of spinal metastases aims to relieve pain, preserve neurological function, maintain the spine’s mechanical stability, and improve overall patient quality of life. Medical treatment can include hormonal therapy for lesions secondary to breast and prostate cancer, chemotherapy, corticosteroids to treat inflammation, bisphosphonates to prevent bone resorption, and analgesia to relieve pain [4, 6]. Recent advances in stereotactic radiosurgery allow for the precise targeting of spinal metastases, delivering effective doses of radiation in cases of limited epidural compression. However, surgical decompression is required for higher-grade ESCC [7].

The surgical removal of spine metastases may be achieved through *en-bloc* resection (wide excision margins to remove the tumor in a single piece), curettage/piecemeal (removal by pieces with no clearly defined margins), or a partial palliative resection for spinal cord decompression [8, 9]. Additionally, percutaneous vertebroplasty and kyphoplasty are minimally invasive vertebral augmentation procedures used to stabilize painful pathologic compression fractures [2, 6]. Although the risk of complications may be high, surgery for spinal metastases provides pain relief and neurological improvement [3]. Polymethylmethacrylate (PMMA) bone cement is commonly used for the reconstruction of the vertebral body post-resection as it is widely available, inexpensive, mechanically strong, and conforms to the shape of the bone when inserted [10]. However, the risk of cancer recurrence remains after a resection. The median time to recurrence for isolated spinal metastatic tumors is 24 months when an *en bloc* resection is performed, with the risk of recurrence increasing when an intralesional curettage is performed [11]. One study determined the local recurrence rate to be 14% in patients who received a percutaneous PMMA vertebroplasty to treat spinal metastases secondary to breast cancer (mean follow-up time of 42 months) [12]. In another study, one in six (17%) patients who underwent an *en bloc* resection experienced a local recurrence. Furthermore, most recurrences were diagnosed 2 years or more after the initial surgery [13].

Studies have investigated bone cement formulations containing antibiotics for the control of local infections in orthopedic applications since the 1970s. PMMA is most often mixed with the antibiotic vancomycin, and clinical outcomes are favourable for infection control [14]. Although these antibiotic PMMA cements are commercially available for clinical use, they are limited in their ability to control the release rate of the drugs and mixing the antibiotic powders directly into the cement can decrease its mechanical properties [15–17]. In the same way antibiotics have been mixed into PMMA for infection control, there is interest in developing PMMA cements loaded with chemotherapy drugs for decreasing the rate of local tumor recurrence in bone related cancers. To date, most studies have mixed chemotherapeutics such as doxorubicin, methotrexate, and cisplatin, directly into the PMMA powder [18, 19]. In these studies, only 5-20% of the drugs mixed into the cement were eluted with most of the release occurring within 24 hours, likely due to the surface burst release effect, since PMMA is non-biodegradable [20].

To increase antibiotic elution from PMMA cement, researchers have incorporated additives including borosilicate glass, nanotubes, mesoporous silica nanoparticles, polycaprolactone (PCL) and polyethylene glycol (PEG). With these additives, an increase in the drug elution occurs [21–25]. However, soluble fillers such as PCL, PEG, or even the drug powders themselves, are porogens that decrease the mechanical strength of PMMA, making it unsuitable for load-bearing applications [23, 26]. One study determined that with the incorporation of 8.15% w/w mesoporous silica nanoparticles into PMMA bone cement, gentamicin elution was significantly enhanced without a negative impact on the cement’s mechanical properties [25]. More recently, a PMMA composite with 15% w/w *γ*-cyclodextrin polymeric microparticles was used to enhance doxorubicin release, with 100% release in 100 days. However, the compressive strength fell below the minimum threshold according to the ASTM F451-16 and ISO 5833 standards [27]. To our knowledge, studies investigating PMMA composites as a chemotherapeutic delivery device are limited. As side effects of systemic chemotherapy administration are common (e.g., doxorubicin-induced cardiotoxicity [28]), such a cement would function as an adjuvant therapeutic device with the potential to bypass negative side effects associated with high doses of systemic chemotherapy administration [29].

There is growing interest in the application of mesoporous silica nanoparticles as drug carriers due to their high surface area, chemical stability, modifiable pore size, and ability to be functionalized [30]. In this study, we developed a PMMA cement functionalized with doxorubicin-loaded nanoparticles for a sustained drug release. We evaluated its mechanical strength and *in vitro* efficacy against MDA-MB-231 breast cancer and C4-2B prostate cancer cell lines in both 2D culture and 3D tumor spheroid cell culture models. Our work is the first to investigate the application of mesoporous silica nanoparticle technology to PMMA bone cement for local chemotherapy drug delivery. Such nanoparticle-functionalized cements could be used to deliver a local dose of chemotherapeutics to lower the risk of recurrence after a spinal tumor resection surgery and improve patient outcomes.

## Materials & Methods

### Nanoparticle doxorubicin loading

Commercially available mesoporous silica nanoparticles (NPs) with a diameter of 500 nm and a pore size of 2 nm were used (Sigma-Aldrich, #805890). NPs were weighed and placed in glass scintillation vials with 1 mL of phosphate-buffered saline (PBS) (Gibco, #10010). The vials were placed in an ultrasonication ice water bath for 2 h to suspend the NPs. Following sonication, doxorubicin hydrochloride (DOX) (Sigma-Aldrich, #44583) solutions prepared in 1mL of PBS were added to the vials to achieve final concentrations of 0, 40, 80, 120, and 240 µM of DOX. The vials were placed on a nutating mixer (fixed speed of 24 rotations per minute) for 24 h at room temperature to allow the drug to be adsorbed into the NPs. The following day, the DOX-NP solutions were transferred to 2 mL Eppendorf tubes and centrifuged at 10,000 rpm for 3 minutes, aggregating the NPs into a pellet. The supernatant was removed and stored to quantify the concentration of unbound DOX. The pellet of DOX-loaded NPs was then dried overnight in a 37°C oven.

### Nanoparticle-cement preparation

To prepare the cement, the NP pellet was crushed and thoroughly mixed into approximately 0.326 g of SmartSet HV bone cement powder (DePuy, #3092040). To the NP-cement powder, 167.2 µL (0.154 g) of liquid methylmethacrylate monomer pre-cooled to 4°C was added and mixed thoroughly for approximately 30 seconds with a spatula. Once the cement achieved a workable consistency, it was mixed on the palm of the hand until it was no longer adhering to the glove, for about 60 seconds. The mixed cement was then placed into a 1 mL BD Tuberculin Slip Tip syringe and was extruded to form a log of cement. A 3D-printed jig was used to cut the malleable cement rod into cylinders of approximately 3 mm in height and 2 mm in diameter. For *in vitro* assays, the cement pods were placed under a UV light overnight for sterilization. Cement with DOX without NPs (Ø NP) was prepared with an equivalent of the 120 µM concentration (240 nmol). The DOX solution was mixed into the cement powder and dried on a hotplate before the liquid fraction was added and the cement was mixed. The various cement compositions are outlined in Table 1.

**Table 1:** Cement formulations

| Sample | Cement Powder (g) | Cement Liquid (mL) | NPs (mg) |  | DOX loaded (nmol) |
| --- | --- | --- | --- | --- | --- |
|  |  |  | 2% w/w | 8% w/w |  |
| Blank | 0.326 | 0.167 | 0 | 0 | 0 |
| ∅ NP | 0.326 | 0.167 | 0 | 0 | 240 |
| 0$\mu\text{M}$ | 0.326 | 0.167 | 10 | 40 | 0 |
| 40$\mu\text{M}$ | 0.326 | 0.167 | 10 | 40 | 80 |
| 80$\mu\text{M}$ | 0.326 | 0.167 | 10 | 40 | 160 |
| 120$\mu\text{M}$ | 0.326 | 0.167 | 10 | 40 | 240 |
| 240$\mu\text{M}$ | 0.326 | 0.167 | 10 | 40 | 480 |

### Drug Release Kinetics

The cylindrical cement pods were weighed, and their dimensions were measured with calipers to the nearest 0.02 mm before being placed into 2 mL Eppendorf tubes with 500 µL of PBS to fully submerge the cement pods. The time of PBS addition was considered the zero timepoint, namely, day 0. After 24 h (day 1), 250 µL of PBS was removed and stored to quantify the concentration of DOX. The volume withdrawn (250 µL) was replenished with fresh PBS to regain the total volume of 500 µL. This was repeated on days 4, 7, 14, 21, and 28. The basis of measuring the concentration from the aliquots of withdrawn PBS relied on the autofluorescence of DOX (excitation: 470 nm, emission: 585 nm). The fluorescence intensity was measured with a TECAN Infinite M200 Pro microplate reader (TECAN, Männedorf, Switzerland) and the concentrations were interpolated from a standard curve. The cumulative DOX concentrations were calculated recursively using the following equation:

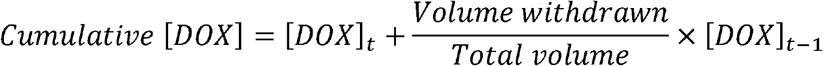

where [DOX]_*t*_ is the concentration of DOX at time *t* and [DOX]_*t*–1_ is the concentration of DOX at time prior to *t* (previous timepoint).

### Mechanical compression testing

To evaluate the mechanical properties of the cement, cylindrical samples were prepared with a diameter of 6mm and a height of 12mm, as per the ASTM F451-16 and ISO 5833:2002 standards. Cement without NPs (blank) and with 8% w/w NPs was mixed and molded into cylinders using a cardboard tube with an inner diameter of 6mm. Once cured, the tubing was removed, and the ends were sanded square to the correct height using 320 grit sandpaper. The dimensions were measured to the nearest 0.02mm using calipers. Uniaxial compression testing was performed on an Instron 1362 (Instron, Norwood, Massachusetts, USA) with a ReNew MTS controller (MTS Systems Corporation, Eden Prairie, Minnesota, USA) and a 20kN load cell at an unconfined compression rate of 20mm/min. The Young’s moduli (moduli of elasticity) were calculated as the slopes of the linear portions of the stress-strain curves. The ultimate compression strengths were determined from the upper yield point loads on the stress-strain curves.

### Scanning electron microscopy

Samples were prepared as above without DOX and cured. They were then mounted in varying orientations onto aluminium stubs using double-sided carbon tape. Samples were then coated with a 4 nm layer of platinum using a ACE600 high resolution sputter coater (Leica Microsystems) before being imaged using an FEI Quanta 450 ESEM (Thermo Fisher Scientific, Saint Laurent, QC).

### Cell lines & in vitro efficacy assays

The cell lines employed in this study were the triple-negative MDA-MB-231 breast cancer cell line and the bone metastatic C4-2B prostate cancer cell line. To obtain DOX dose-response curves for these cell lines, 20,000 cells in the log phase of growth were seeded in 48-well tissue culture plates. MDA-MB-231 and C4-2B cells were seeded in DMEM (Gibco, #12430062) and RPMI 1640 (Gibco, #11835055), respectively, supplemented with 10% heat-inactivated fetal bovine serum (FBS) (Gibco, #12483020) and 1% penicillin-streptomycin (Gibco, #15140122). The following day, the cell medium was removed and replaced with a low-serum (1% FBS) media containing the following concentrations of DOX: 0 (PBS vehicle control), 0.001, 0.01, 0.05, 0.1, 0.5, 1, 5, 10, and 100 µM. AlamarBlue (Invitrogen, #DAL1100) resazurin reduction assays were performed after 3 and 7 days of DOX treatment to determine metabolic activity compared to control. Briefly, cell medium was removed and a 10% AlamarBlue solution in low-serum medium was added and the plates were incubated at 37°C for 4h. The resorufin fluorescence was quantified using the TECAN fluorescence microplate reader (excitation: 560nm, emission: 590nm).

To determine the *in vitro* efficacy of the DOX cement against tumor cells in a 2D monolayer culture, the cement constructs loaded with various concentrations of DOX were incubated with cultures of 20,000 cells in 48-well tissue culture plates, as described above. The cement pods were submerged directly into the wells with low-serum culture medium. AlamarBlue assays were performed after 1, 4, and 7 days of cement treatment to determine metabolic activity. Live/Dead (Invitrogen, #L3224) cell viability/cytotoxicity stains were applied according to manufacturer instructions. Briefly, cell medium was removed and 100µL of Live/Dead solution in PBS was added to the wells and incubated at 37°C for 15 minutes before being imaged with an EVOS M5000 imaging system (Invitrogen, #AMF5000).

To assess the efficacy of the cement in 3D *in vitro* culture, MDA-MB-231 and C4-2B spheroids were formed by placing 20,000 cells in Nunclon Sphera round-bottom 96-well plates (Thermo Fisher Scientific, #174925) and centrifuging at 290xG for 3 minutes to form a pellet (day 0). On day 1, MDA-MB-231 pellets received 100µL of a 4µg/mL solution of ready-to-use bovine collagen I (Sigma-Aldrich, #5074) for a final concentration of 2µg/mL and were centrifuged at 100xG for 3 minutes. C4-2B spheroids were ready on day 4 and MDA-MB-231 spheroids were ready on day 6. Once ready, the cell medium was discarded, and spheroids were coated with a 1:1 solution of low-serum cell culture medium and collagen I gel for a final collagen concentration of 2.5mg/mL. Spheroids were considered ready once they were compact enough to tolerate the addition of the 2.5mg/mL collagen solution without breaking apart. The plates were incubated at 37°C for 90 minutes for the gelation of the collagen. Following the gelation, the wells were topped with fresh low-serum cell culture medium. The following day, the spheroids in collagen gel were transferred to non-adherent 48-well plates containing low-serum cell culture medium, and the cement pods were juxtaposed in the wells. A 2µM concentration of free DOX in cell medium served as a positive control. AlamarBlue assays were performed after 1, 4, and 7 days of cement treatment to determine metabolic activity, with a modified incubation time of 6h. Live/Dead and DAPI (Invitrogen, #R37606) stains were applied according to manufacturer instructions. Fluorescence and phase contract images of the pellets were taken to visualize pellet size and cell migration into the collagen matrix. Area quantification was performed with ImageJ software (NIH, Maryland, USA).

### Statistics

GraphPad Prism Version 9 (GraphPad Software, San Diego, California, USA) was used for statistical analyses. Student’s *t* tests were used to test for significant differences in mechanical properties between cement groups (blank vs. 8% w/w NP). DOX release data were fitted with two-phase exponential association functions. The compounded standard error of the mean (SEM) was determined for DOX release data due to the propagation of error from repeated measurements of the same samples at various timepoints. Dose-response data were fitted using sigmoidal functions and relative IC_50_ values were obtained with their 95% confidence intervals. Two-way ANOVAs with Tukey’s correction were performed to test for significance between *in vitro* DOX cement treatments at multiple timepoints. Experiments were run in triplicate and performed thrice for an *N* of 3, unless otherwise indicated. Significance was set at an alpha of 0.05. Where shown in-text, data are represented as mean ± SD. Graphical error bars are represented as SEM and ns indicates *p* > 0.05, * indicates *p* < 0.05, ** indicates *p* < 0.01, *** indicates *p* < 0.001, and **** indicates *p* < 0.0001.

## Results

### Doxorubicin-Nanoparticle Cement & Release Kinetics

Mesoporous silica NPs were loaded with DOX and mixed into PMMA bone cement. The cement was shaped into cylindrical pods with a diameter of 2.23 ± 0.08mm and a height of 3.13 ± 0.07mm. The addition of NPs to the cement created a network and altered the microstructure within the cement, creating pockets of NPs encapsulated by the cement (Figure 1). The polymer beads in the cement powder interconnected with the polymerization of the methylmethacrylate and incorporated the NPs, fixing them in place. The cement containing 8% w/w NPs displayed a higher degree of connectivity between NPs compared to cement containing 2% w/w NPs.

**Figure 1:**
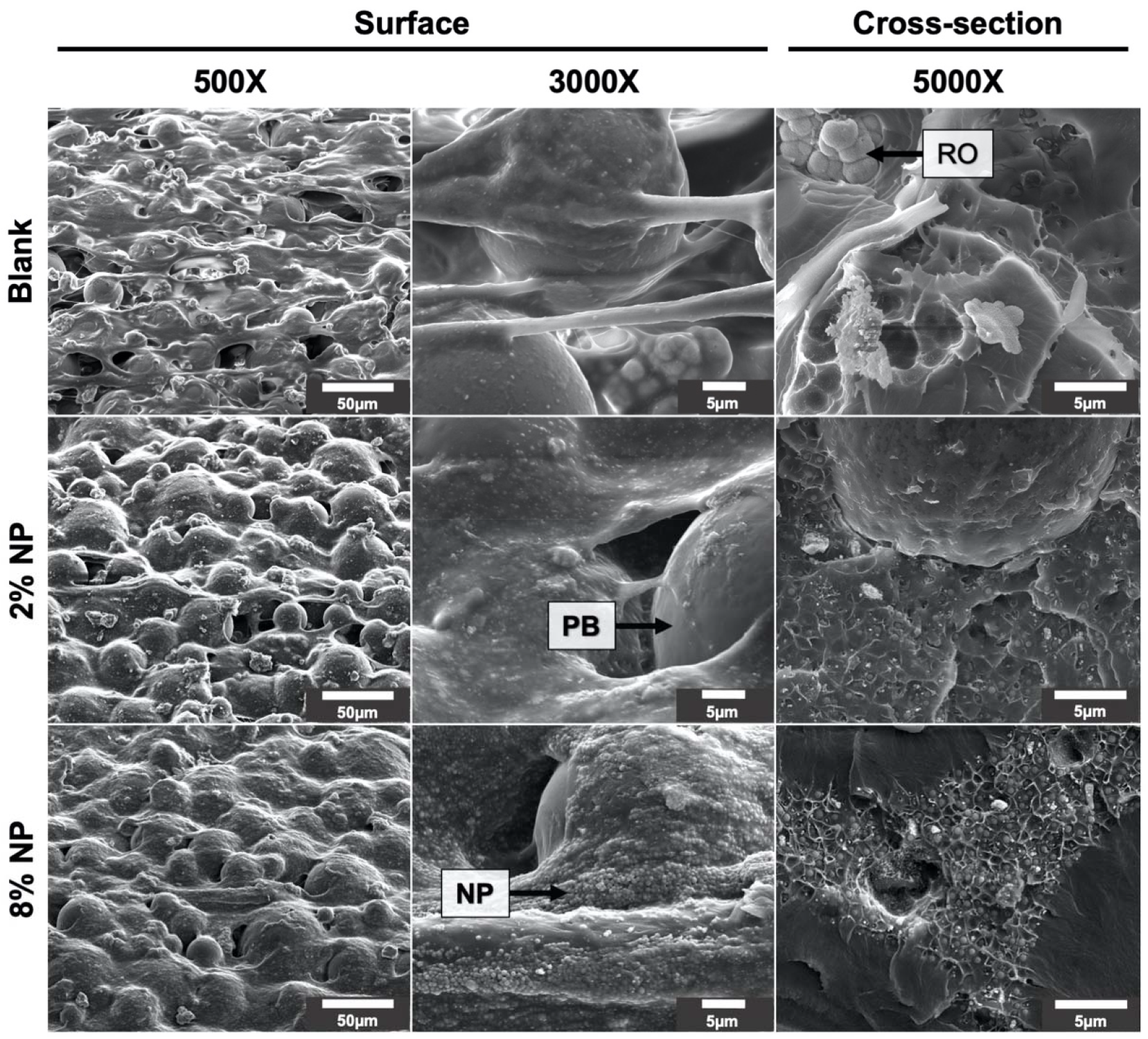
Surface and cross-sectional scanning electron microscope images of cement mixed without NPs (blank) and with 2% and 8% w/w NPs. RO: Radioopacifier crystals (zirconium dioxide; ZrO_2_). PB: PMMA polymer beads. NP: nanoparticles.

The mechanical properties of cement were evaluated with the addition of nanoparticles. Cylindrical samples of blank cement were prepared with a diameter of 6.03 ± 0.03mm and a height of 12.00 ± 0.05mm, and cement containing 8% w/w NPs was prepared with a diameter of 6.12 ± 0.06 mm and a height of 11.98 ± 0.02 mm. Representative stress-strain curves are shown in Figure 2A. The Young’s modulus for blank cement was determined to be 2983 ± 99 MPa and 3024 ± 90 MPa for 8% w/w NP cement, revealing no significant difference between the groups (Figure 2B, *p* > 0.05). Furthermore, the ultimate compressive strength of blank cement was determined to be 120 ± 4 MPa and 116 ± 7 MPa for 8% w/w NP cement, with no significant difference between the groups (Figure 2C, *p* > 0.05).

**Figure 2:**
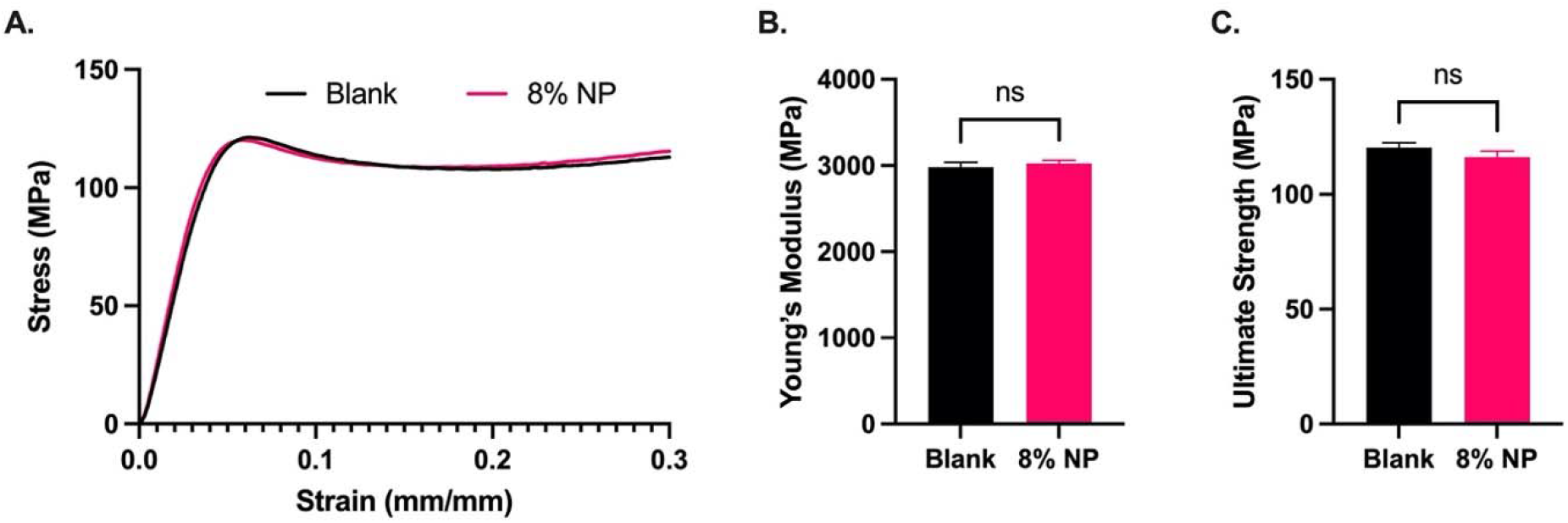
(**A**) Stress-strain curves, (**B**) Young’s moduli, and (**C**) ultimate compression strengths of cement containing no NPs (blank; *N* = 3) and 8% w/w NPs (*N* = 5). Error bars represent SEM.

The release kinetics of DOX were evaluated in cement containing 2% and 8% w/w NPs, loaded with various amounts of DOX. The NPs retained between 77% and 93% of the DOX in the loading solution. At a given quantity of NPs, the proportion retained increased with increasing concentrations of DOX in the loading solution (Figure 3A). Additionally, 40mg of NPs were able to retain approximately 5% more DOX compared to 10mg at a given loading concentration (*p* < 0.05). When cement was mixed with 240nmol of DOX without NPs, release was limited, and a plateau was reached by day 4 (Figure 3B). With the addition of NPs, the release of DOX was facilitated with a significant increase in the cumulative DOX release at day 4 (*p* < 0.05). Furthermore, cement containing 8% w/w NPs had a significantly higher cumulative release on day 4 compared to the equivalent cement containing 2% w/w NPs (*p* < 0.05). By day 28, the cement with 2% and 8% w/w NPs had released 3.1 and 16.7 times more DOX, respectively, compared to the cement without NPs.

**Figure 3:**
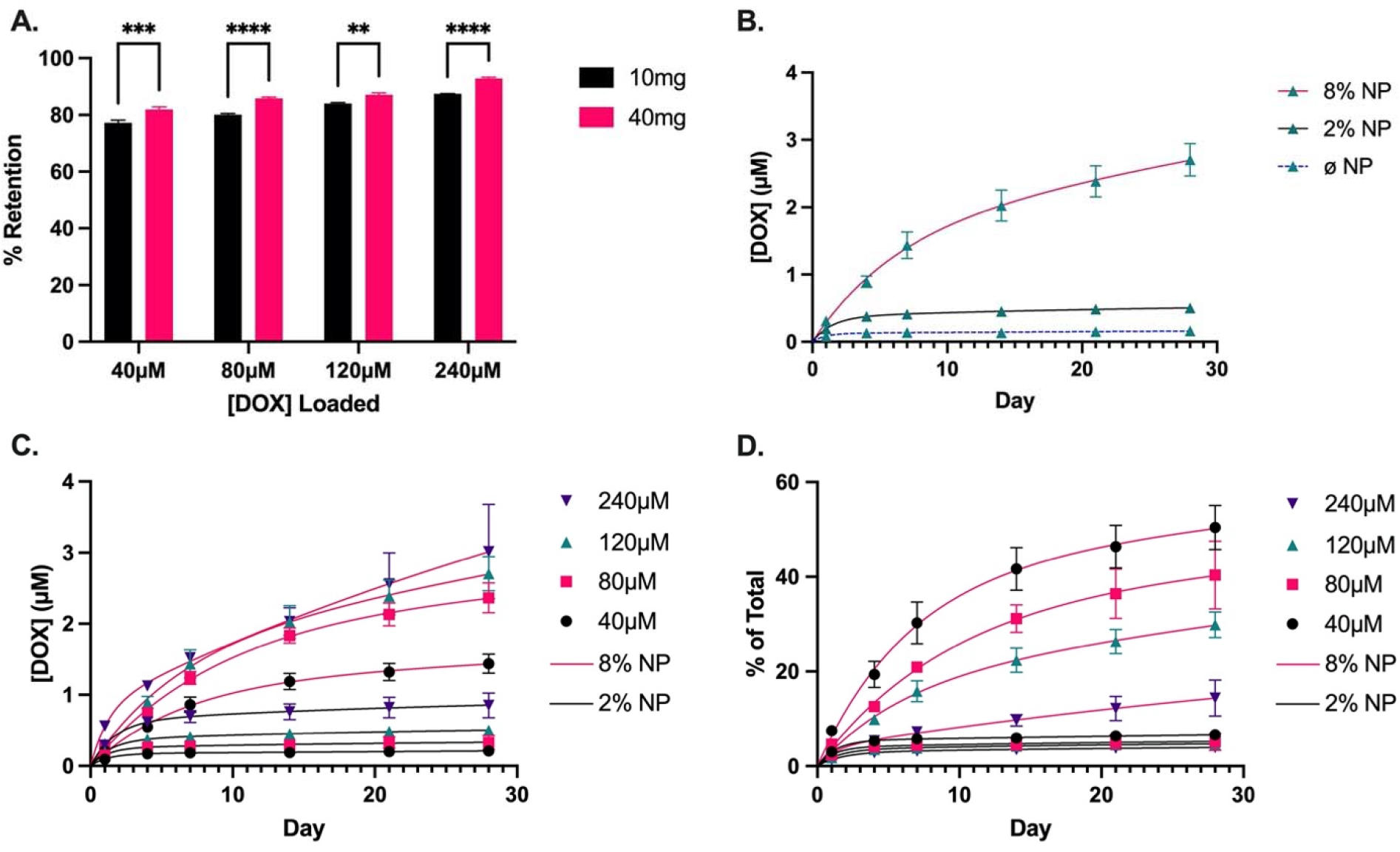
(**A**) Proportion of DOX retained by 10 and 40mg of NPs at various loading concentrations of DOX, corresponding to 2% and 8% NP w/w mixed into the cement, respectively; (**B**) DOX release from cement containing no NPs (Ø NP; *N* = 2), 2% and 8% w/w NPs loaded with an equivalent of 120µM of DOX; (**C**) Cumulative concentration and (**D**) proportional release of DOX over 28 days from cement containing 2% and 8% w/w NPs loaded with 40, 80, 120, and 240µM of DOX. Solid line: 2% w/w NP. Dashed line: 8% w/w NP. *N* = 3. Error bars represent SEM in (**A**) and compounded SEM for DOX release profiles in (**B**–**D**).

The cumulative release of DOX was higher for the cement containing 8% w/w NPs compared to 2% w/w NPs at all concentrations of DOX as of day 7 (Figure 3C, *p* < 0.05). Within each cement group (2% and 8%), the cumulative release was higher for the samples loaded with higher concentrations of DOX (*p* < 0.05). Similarly, the proportional release of DOX was higher for the cement containing 8% w/w NPs compared to 2% w/w NPs (Figure 3D). However, there was no significant difference in the proportional release between the various concentrations for cement containing 2% w/w NPs, with an average proportional release rate of 0.037% per day over days 14–28. Conversely, there was a significant difference in the proportional release rates of DOX within the 8% w/w NP cement group. The cement containing NPs loaded with lower concentrations achieved a higher proportional release, as high as 50% in 28 days. The average proportional DOX release rate for 8% w/w NP cement was 0.54% per day over days 14–28, or approximately 15 times higher compared to 2% w/w NP cement. Assuming a constant diffusion coefficient (*D*) for finite cylinders, the release kinetics are governed by the following:

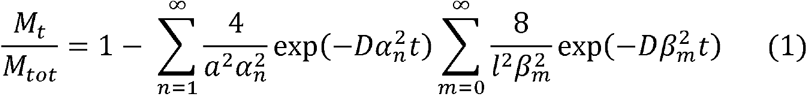

where *M*_*t*_ is the mass of DOX released at time *t, M*_*tot*_ is the total mass of DOX in the sample, *a* is the radius and *l* is the height of the cylinder, *aα*_*n*_ is the n^th^ zero of the zero-order Bessel function of the first kind where *J*_0_(*aα*_*n*_) = 0, and 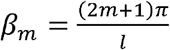. At later stages of release, only the first term in Eq. 1 will contribute and we obtain:

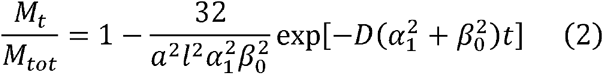

which gives us the following expression:

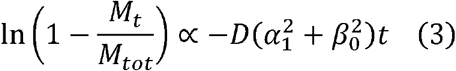

Thus, by plotting ln(1 − *M*_*t*_/*M*_*tot*_) versus time (Figure 4) we are able to calculate the slope of the linear region to obtain the average diffusion coefficients for each cement composition:

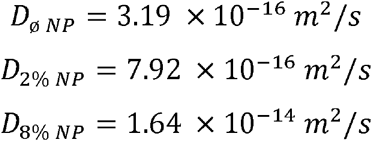

**Figure 4:**
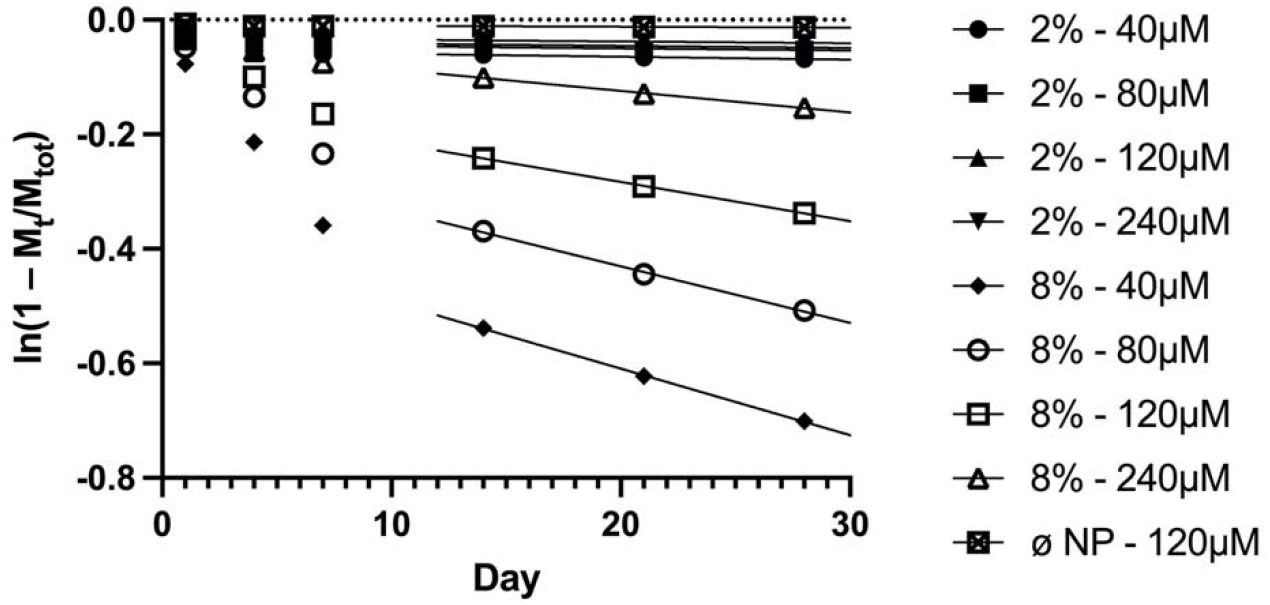
Plot of ln(1 – *M*_*t*_/*M*_*tot*_) versus time, where *M*_*t*_ is the mass of DOX released at time *t* and *M*_*tot*_ is the total mass of DOX in the sample.

### 2D *In Vitro* Efficacy Assay

Both MDA-MB-231 and C4-2B cells treated with DOX for 7 days had higher IC_50_ values compared to cells treated for 3 days (*p* < 0.05). The IC_50_ curves are shown in Figure 5. Table 2 provides the calculated IC_50_ values. These IC_50_ values guided the selection of the loading concentrations of DOX for the NPs to achieve the release of effective drug doses.

**Table 2:** IC_50_ values for DOX against MDA-MB-231 and C4-2B cell lines at D3 and D7, with 95% confidence intervals (95% CI).

| Cell Line | $IC_{50}$ – D3 (μM) | $IC_{50}$ – D7 (μM) |
| --- | --- | --- |
| MDA-MB-231 | 0.14 (95% CI: 0.09, 0.21) | 0.48 (95% CI: 0.41, 0.56) |
| C4-2B | 0.07 (95% CI: 0.05, 0.09) | 0.32 (95% CI: 0.26, 0.40) |

**Figure 5:**
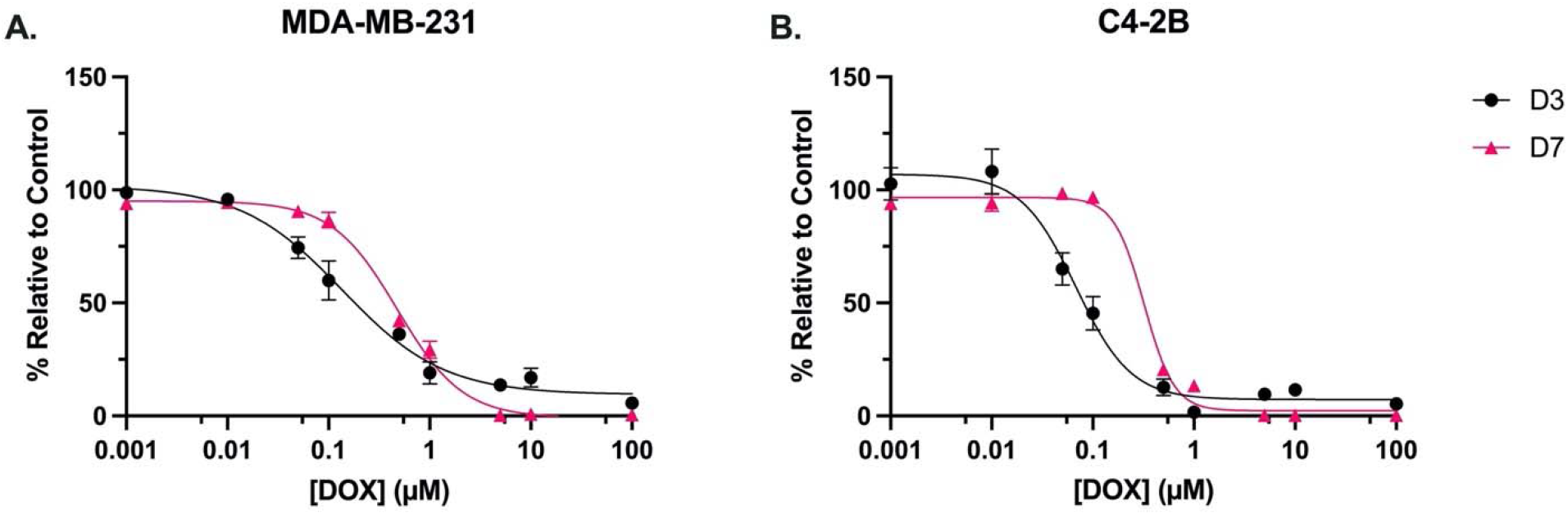
Dose-response curves evaluating the metabolic activity of (**A**) MDA-MB-231 and (**B**) C4-2B cells treated with DOX for 3 days (D3) and 7 days (D7). *N* = 3. Error bars represent SEM.

The *in vitro* efficacy of the DOX-loaded NP cement was first evaluated in monolayer cultures of MDA-MB-231 and C4-2B cells. At day 1, there was no significant difference between the treatments for both cell lines. After 4 and 7 days of treatment, the metabolic activity of MDA-MB-231 cells was significantly reduced with the DOX cement formulations compared to the cement containing NPs without DOX (0µM) (Figure 6A, *p* < 0.05). Furthermore, on day 4, the cement containing NPs loaded with 120µM of DOX was more effective than the 40µM formulation (*p* < 0.05). There was no significant difference between the metabolic activities of MDA-MB-231 cells treated with and without NPs (blank). C4-2B cells treated with 0µM cement had a lower metabolic activity on day 4 compared to blank cement (Figure 6B, *p* < 0.05). This difference was no longer significant at day 7. After 4 and 7 days of treatment, the metabolic activity of C4-2B cells was significantly reduced with the DOX cement formulations compared to the 0µM cement (*p* < 0.05). Representative images of Live/Dead staining are shown in Figure 7. For both cell lines, a reduction in cell proliferation and viability on day 7 compared to day 1 is observed in cells treated with DOX cement.

**Figure 6:**
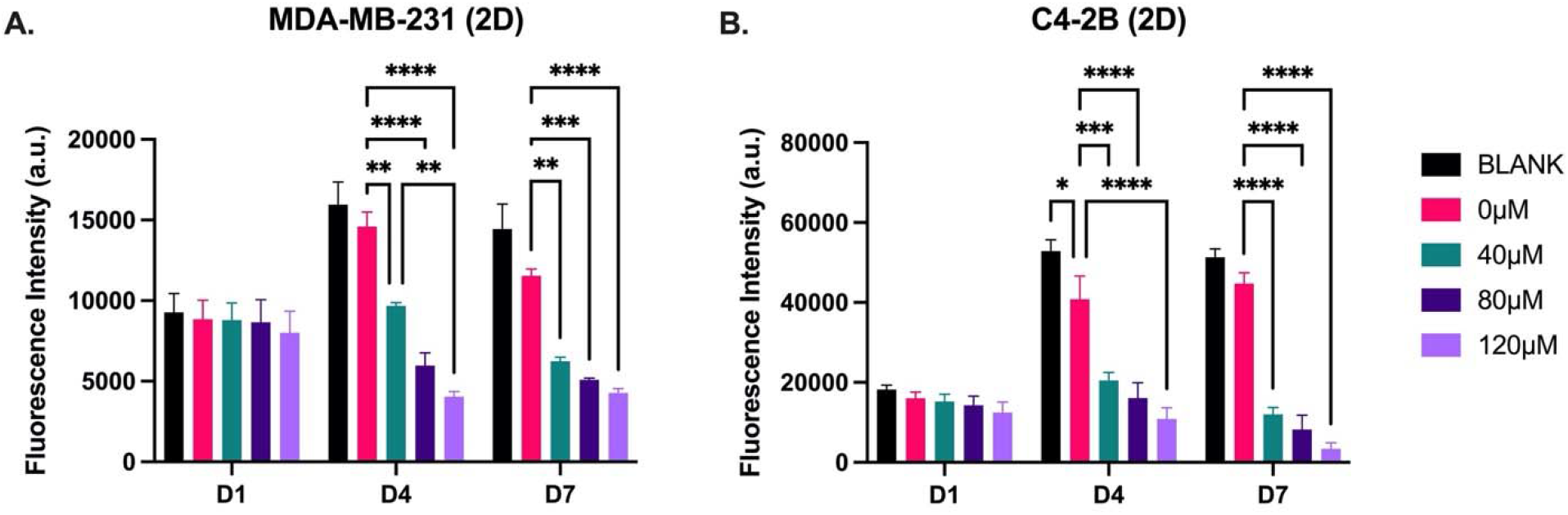
Metabolic activity of 2D cultures of (**A**) MDA-MB-231 and (**B**) C4-2B cell lines treated with cement constructs for 1 day (D1), 4 days (D4), and 7 days (D7). Blank: cement without NPs. 0µM: NP cement without DOX. *N* = 3. Error bars represent SEM.

**Figure 7:**
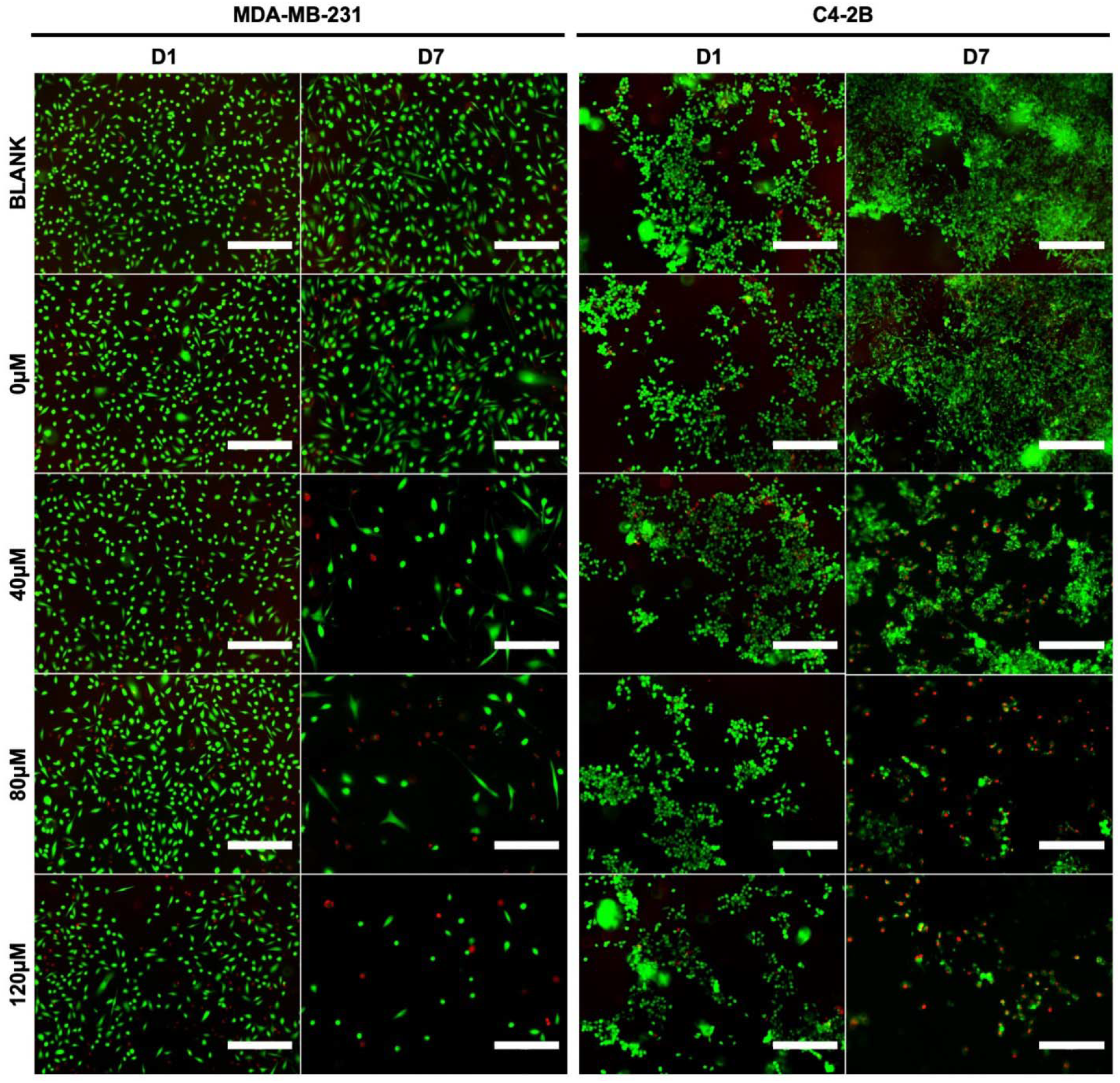
Representative Live/Dead staining images of MDA-MB-231 and C4-2B cells at day 1 (D1) and day 7 (D7) of treatment with cement constructs. Green: alive; red: dead. Scale bar = 300µm.

### 3D *In Vitro* Metabolic Activity Assay & Pellet Size Quantification

To assess the *in vitro* efficacy of the DOX cement constructs in a more physiological cell culture system, metabolic activity, spheroid growth and cell outgrowth was assessed in 3D collagen-coated spheroid cultures. This method allowed for the concurrent evaluation of metabolic activity and cell migration. There was no significant difference in terms of metabolic activity and cell migration between treatments on day 1 for both MDA-MB-231 and C4-2B spheroids. On days 4 and 7, there was a significance reduction in metabolic activity of MDA-MB-231 spheroids treated with DOX cement, compared with the 0µM cement (Figure 8A, *p* < 0.05). On day 7, MDA-MB-231 spheroids treated with the 0µM displayed lower metabolic activity compared to the blank cement (*p* < 0.05). The activity of C4-2B spheroids was significantly reduced with the 120µM composition on day 4, and with the 240µM composition on day 7 (*p* < 0.05). A high degree of cell migration was observed in the collagen-coated MDA-MB-231 spheroids from day 1 to 7 for the blank and 0µM cement treatments (Figure 9). The area of migrated cells was significantly reduced on days 4 and 7 with the DOX cement, compared to the 0µM cement (*p* < 0.05). The area of cell migration was lower in the 0µM cement compared to the blank cement on day 7 (*p* < 0.05). Furthermore, the MDA-MB-231 spheroids displayed necrotic tumor cores on day 7 for all treatments and the spheroids treated with DOX appear to have gradually imploded, showing necrotic rings around the inner core (Figure 9). C4-2B spheroids did not display a high degree of cell migration but did show differences in overall spheroid size and compaction (Figure 10). By day 4, the 240µM cement formulation significantly reduced C4-2B spheroid size compared to the 0µM cement (*p* < 0.05). By day 7, both the 120µM and 240µM formulations were effective at inhibiting the growth of C4-2B spheroids (*p* < 0.05). The Live/Dead staining of the C4-2B spheroids treated with the DOX cement show the highest degree of cell death in the spheroid core, with a darker appearance overall with phase contrast imaging (Figure 10).

**Figure 8:**
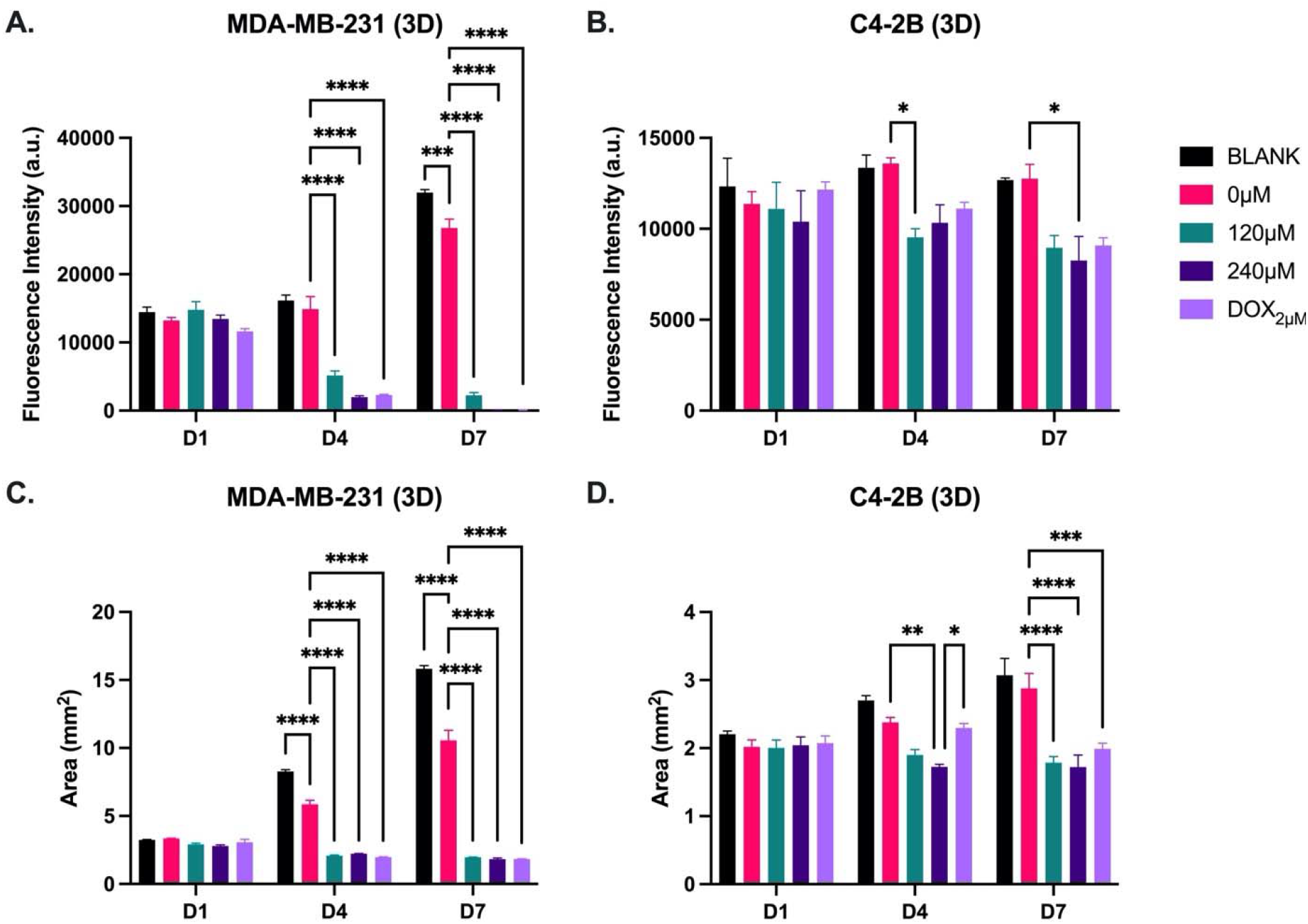
Metabolic activity and area of 3D spheroid cultures of (**A, C**) MDA-MB-231 and (**B, D**) C4-2B cell lines treated with cement constructs for 1 day (D1), 4 days (D4), and 7 days (D7). DOX_2µM_: free 2µM DOX treatment. *N* = 3. Error bars represent SEM.

**Figure 9:**
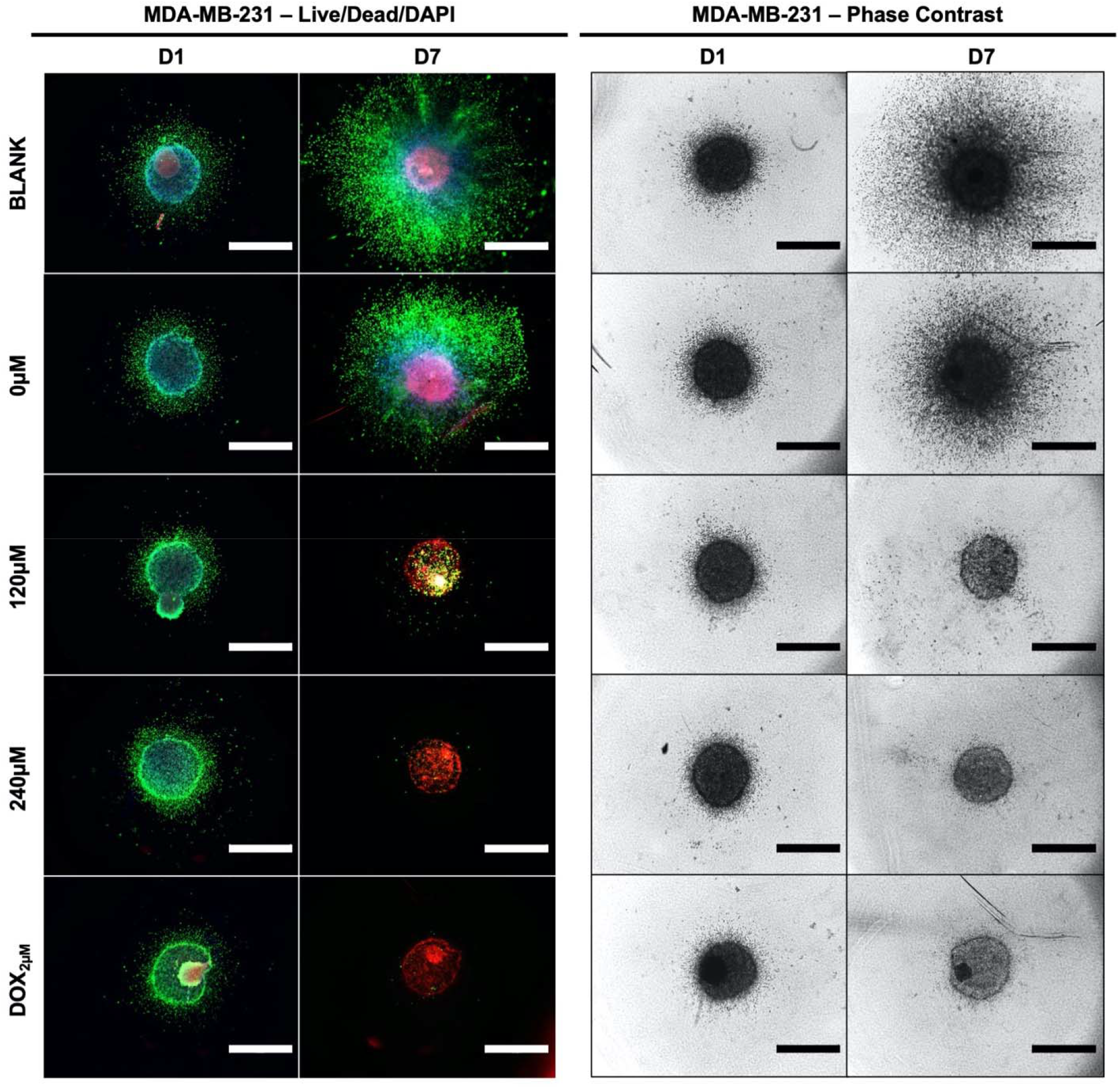
Representative Live/Dead/DAPI staining and phase contrast images of MDA-MB-231 cells at day 1 (D1) and day 7 (D7) of treatment with cement constructs. Green: alive; red: dead; blue: cell nuclei. Scale bar = 1250µm.

**Figure 10:**
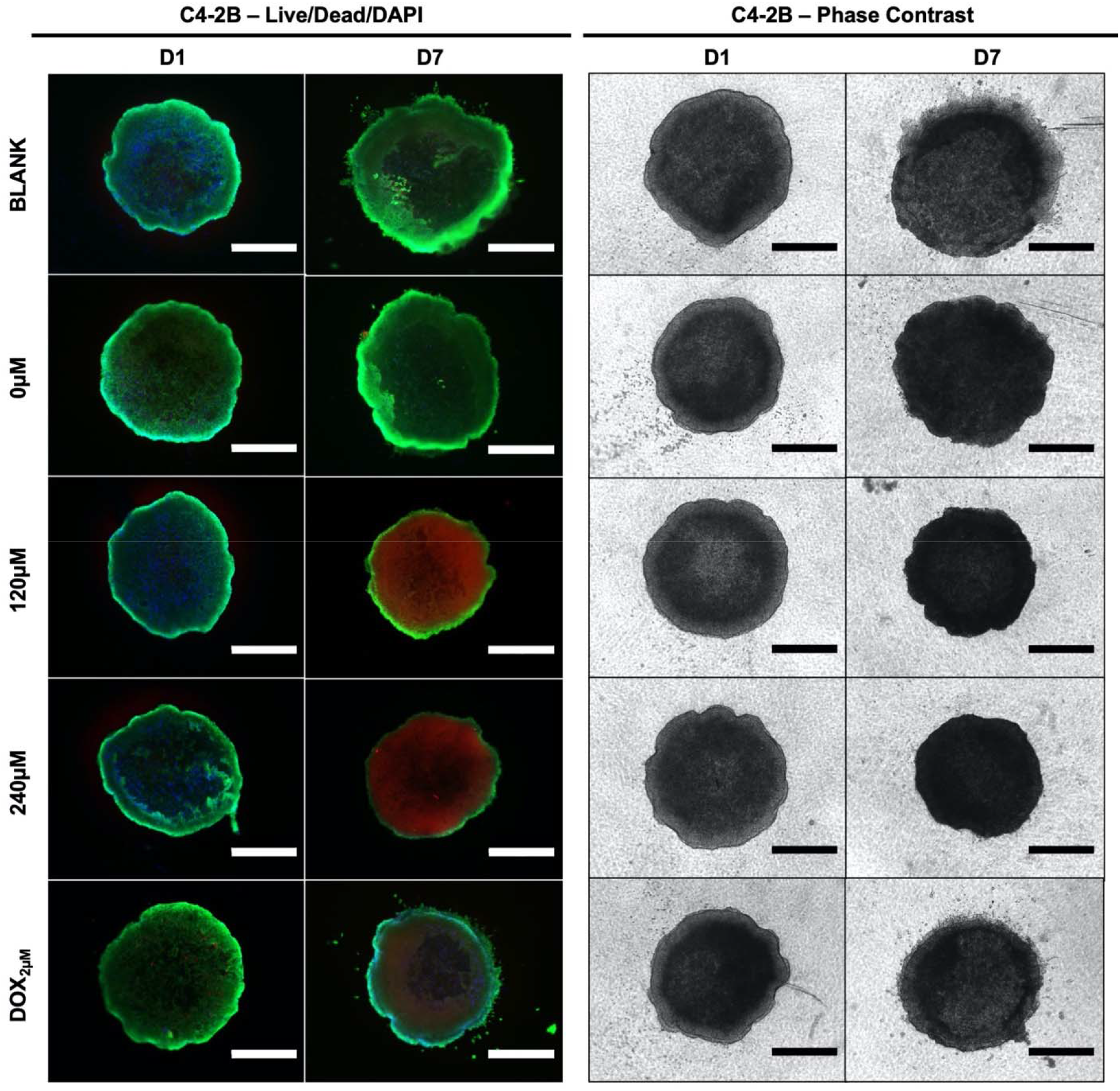
Representative Live/Dead/DAPI staining and phase contrast images of C4-2B cells at day 1 (D1) and day 7 (D7) of treatment with cement constructs. Green: alive; red: dead; blue: cell nuclei. Scale bar = 750µm.

### 2D vs. 3D *In Vitro* Efficacy

On day 7 of treatment, MDA-MB-231 cells treated with control cement (0µM) in 3D culture had a 2.3-fold higher metabolic activity compared to cells in 2D culture (Figure 11A, *p* < 0.05). There was no significant difference in MDA-MB-231 metabolic activity between 2D and 3D culture when treated with DOX cement. However, after a 7-day treatment with DOX cement, the activity of MDA-MB-231 cells was reduced by 63% compared to control cement in 2D culture and by 92% in 3D culture, demonstrating an increased DOX efficacy in 3D culture against MDA-MB-231 cells. Conversely, C4-2B cells in 3D culture exhibited a 3.5-fold lower metabolic activity after 7 days compared to 2D culture (Figure 11B, *p* < 0.05). There was no significant difference in C4-2B metabolic activity between 2D and 3D culture when treated with DOX cement for 7 days. However, after a 7-day treatment with DOX cement, the activity of C4-2B cells was reduced by 92% compared to control cement in 2D culture and by 30% in 3D culture, demonstrating a reduced DOX efficacy in 3D culture against the C4-2B cell line.

**Figure 11:**
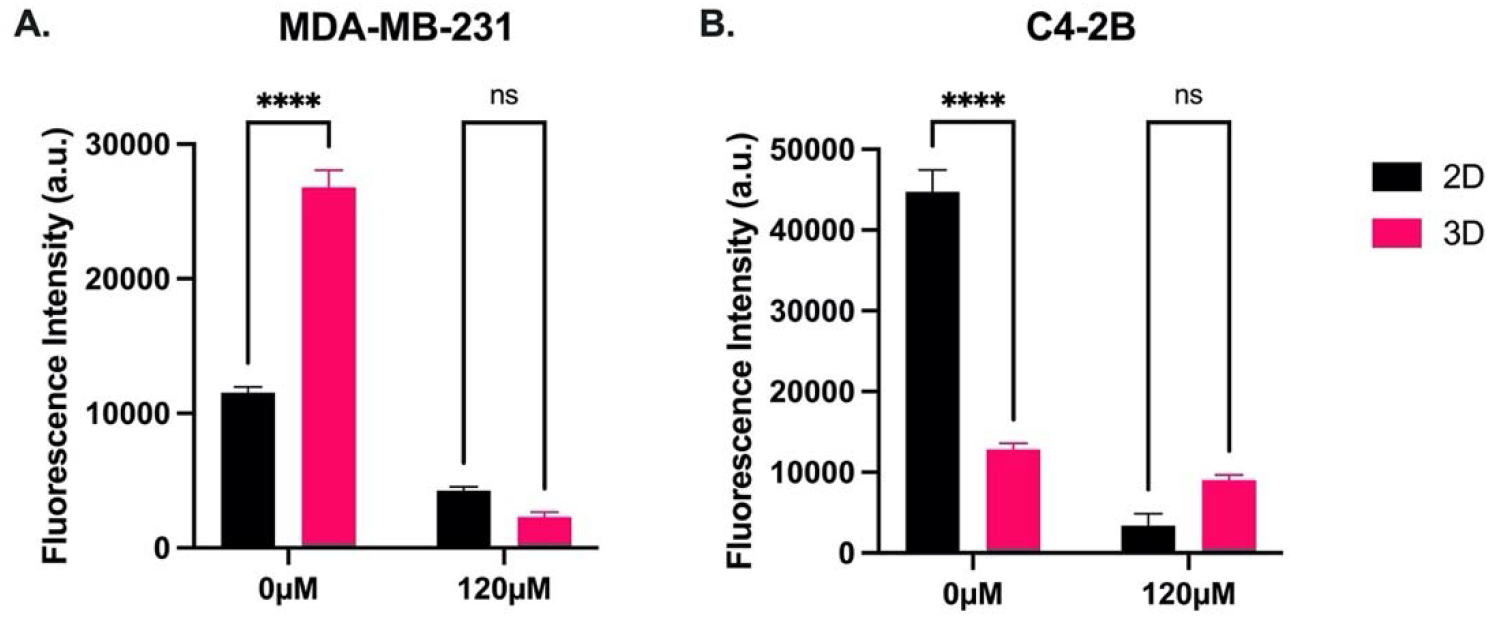
Comparison of the metabolic activity of (**A**) MDA-MB-231 and (**B**) C4-2B cells in 2D and 3D culture treated for 7 days with control cement (0µM) and cement containing NPs loaded with 120µM DOX. *N* = 3. Error bars represent SEM.

## Discussion & Conclusion

The development of a bone cement with both mechanical integrity and controlled chemotherapeutic release is desirable for preventing local tumor recurrence after bone tumor resection surgeries. In the present study, we investigated the addition of doxorubicin-loaded mesoporous silica nanoparticles to PMMA bone cement. Specifically, we evaluated the compressive strength with the addition of nanoparticles, the doxorubicin release profiles as a function of loaded drug concentrations and nanoparticle content within the cement. We then assessed the *in vitro* efficacy of the cement in 2D and 3D cell cultures using breast and prostate cancer cell lines and compared drug sensitivities between 2D and 3D culture models.

With the addition of NPs to PMMA cement, SEM imaging displayed a change in the microstructure of the cement. With a higher content of NPs in the cement, a higher degree of connectivity was observed between the NPs, leading to the formation of a nanonetwork within the cement. PMMA cements functionalized with mesoporous silica NPs require a certain concentration of NPs for the formation of a network that significantly improves drug elution [25]. Based on our drug elution profiles, the 2% NP w/w formulation only improved elution profiles slightly compared to the cement without NPs, with minor increases with increasing loaded concentration of DOX. With the 8% w/w NP composition, the elution rates had a higher dependence on the loaded concentration of the drug with a significantly improved elution rate. This indicates that a sufficient network had been formed to allow for the DOX to diffuse out from not only the cement’s surface, but from within the cement through the interconnected pores of the NPs. Slane *et al*. investigated the mechanical effects of the addition of mesoporous silica NP content in PMMA cement up to a content of 5% w/w. They determined that with increasing NP content, there was an increase in the flexural modulus, compressive strength, and compressive modulus. However, they observed a decrease in flexural strength, fracture toughness, and work to fracture [31]. We found that the addition of 8% NPs w/w to PMMA did not impact the ultimate compressive strength of the cement, nor the Young’s (compressive) modulus, as the stress-strain curves were nearly identical. Similarly, Letchmanan *et al*. demonstrated that the addition of mesoporous silica nanoparticles at 8.15% w/w into PMMA bone cement had negligeable effects on the compressive strength and flexural modulus even after 6 months of aging in PBS, and significantly increased the release of the loaded antibiotic, gentamicin [25]. This demonstrates the feasibility of the NP cement for load-bearing applications, such as in the case of the spine. PMMA cements can be modified with additives to increase their flexural strength and fracture toughness. Khaled *et al*. utilized nanostructured titania fibers and determined that the addition of 1% w/w to PMMA significantly increased its flexural strength, flexural modulus, and fracture toughness [32]. Studies evaluating the mechanical properties of multi-functionalized PMMA cements are necessary to develop a cement that can deliver long-term effective drug doses without compromising mechanical integrity.

It is thought that with the addition of NPs the enhanced drug release is achieved via the creation of a porous nanonetwork in the cement [25, 33, 34]. The mesoporous NPs themselves can hold large amounts of drugs within their pores due to the high surface area. As we increased the concentration of DOX in the loading solution for the NPs, the percentage of the DOX retained by the NPs also continued to increase. This indicates that the NPs had not yet been saturated with the drug at these concentrations. Furthermore, there was only a 5% increase in retention despite a 4-fold increase in the quantity of NPs. This demonstrates the exceptionally high capacity of mesoporous NPs to hold compounds within their pores. Shen *et al*. determined that mesoporous NPs with a diameter of 120nm and a pore size of 5.2nm were capable of retaining DOX at a concentration of 306 mg/g of NPs [35], well beyond what we used in this study. The addition of NPs was required for the cement to release effectives doses of DOX in a sustained manner. Without NPs, a plateau was reached relatively quickly in the release profile. When a higher concentration of DOX was used to load the NPs, they retained more of it and were also able to release it more. With a higher NP content in the cement, the diffusion of DOX was greater for all concentrations. The 2% NP w/w cement loaded with 240µM released less DOX then the 8% NP w/w cement loaded with 40µM, despite being loaded with a six-fold higher DOX concentration. This indicates that the drug elution rate from PMMA is largely determined by the NP content within the cement. The proportion of DOX released from the cement was highest for the cement containing the NPs loaded with the lowest concentration of DOX. This is likely due to the DOX closer to the proximity of the surface being eluted the quickest, and the diffusion from the center being the most difficult [36]. Given a 50% release of DOX in 28 days from the 8% NP w/w cement with the lowest DOX concentration, it is reasonable to postulate that the NP cements with higher amounts of DOX would simply take longer to liberate more DOX, allowing for a prolonged release. It is worth noting that within both 2% and 8% w/w NP cement groups, the diffusion coefficients decreased slightly with increasing loading concentrations. This may be due to the imperfect sink conditions in the method for determining the release kinetics. Nonetheless, the addition of NPs to the cement confers a greater mass diffusivity, with over a 50-fold increase with 8% w/w NPs in the cement compared to without, and over a 20-fold increase compared to 2% w/w NPs.

Consistent with our findings, PMMA with 10.19% NPs w/w loaded with DOX released 60% of the drug in a period of 60 days, whereas the cement without NPs plateaued at 5% of the loaded drug on the first day followed by negligeable release thereafter [34]. To increase the drug releases rates, Li *et al*. functionalized mesoporous NPs with poly(acrylic acid). As a result, the NP drug release profiles were pH sensitive. The functionalized NPs released 70% of DOX in 6 days at pH 5.5, and 25.3% at pH 7.4 [37]. As tumor microenvironments are acidic (pH: 6.5-6.9) due to the high glycolytic rate of tumor cells [38, 39], mixing functionalized NPs such as these into PMMA cement would allow for a pH-controlled drug release, allowing for a tumor-responsive cement. Future work may evaluate the release profiles in terms of nanoparticle diameter, pore size, and functionalization.

To determine the optimal concentration ranges for DOX-NP loading, dose-response experiments were conducted on monolayer cultures of MDA-MB-231 and C4-2B cell lines. We determined DOX IC_50_ values for 3-day (72h) incubation times for MDA-MB-231 and C4-2B cell lines, shown in Table 2. These values were comparable to those in the literature [40, 41]. Additionally, we report the 7-day (168h) IC_50_ values to evaluate drug resistance. As seen in Figure 5, both cell lines display higher cell metabolic activities at the mid-range doses when treated with DOX for 7-days, indicating that the cells are more resistant. Han *et al*. observed that the number of MDA-MB-231 cells exposed to mid-range DOX concentrations had increased after 8 days of treatment, demonstrating their ability to proliferate in a tolerable DOX concentration range [42]. They concluded that the DOX resistance is associated with epigenetic alterations in histone deacetylase, and that suberoylanilide hydroxamic acid (SAHA; a histone deacetylase inhibitor) restored DOX sensitivity [42]. This is an important consideration for a cement intended for a sustained drug release profile, as the dose must be high enough to avoid chemoresistance. Agents such as SAHA can be incorporated into the cement to increase the sensitivity of the tumor cells to the chemotherapy drug. Tonak *et al*. combined SAHA with valproic acid mixed directly into PMMA for the local treatment of osteosarcoma [43]. Similarly, Koto *et al*. directly incorporated the third-generation bisphosphonate, zoledronic acid, into PMMA cement and the resulting cement exhibited antitumor effects against multiple malignant tumor cell types without systemic toxicity *in vivo* [44].

When incubated with both 2D and 3D cultures of MDA-MB-231 and C4-2B cell lines, a time– and dose-dependent inhibition of the cancer cells was observed with the DOX-NP cement formulations. In 2D culture, the metabolic activities of both cell lines were inhibited with all DOX-NP cement formulations as of day 4 and maintained at day 7. On day 4 of treatment, C4-2B cells showed a lower metabolic activity when treated with the 0µM NP cement, compared to the blank cement. However, on day 7 this difference was no longer significant. It is possible that the addition of the NPs to the cement facilitated the release of the methylmethacrylate (MMA) monomer in the cement. Unpolymerized MMA leakage from PMMA has been demonstrated to have cytotoxic effects on human cells, and PMMA cement has been shown to have a cytotoxic effect against metastatic spine cells [45–47]. For both cell lines, Live/Dead imaging from 2D culture displays a reduction in the density of living cells and an increase in dead cells with the DOX-NP cement treatment, as compared to the 0µM and blank controls. Most studies have been conducted with chemotherapy drug powders mixed directly into PMMA. Furthermore, the method used to test the cement *in vitro* varies between studies. Many studies have mixed methotrexate, doxorubicin, and cisplatin directly into PMMA and incubated the cement samples with cell medium, and then treated cells with the medium containing the drugs eluted from the cement. As the release kinetics reach their peak quickly due to a burst release, efficacy against cancer cells decreases over time [19, 48, 49]. Cyphert *et al*. recently used this indirect incubation method to test their DOX-loaded polymeric *γ*-cyclodextrin microparticle PMMA cement [27]. They determined that PMMA loaded with 15% w/w DOX-microparticles had consistent cytotoxic effects over time, compared to cement with DOX mixed directly into it. This can be attributed to the more consistent release of the drug from the composite cement, relative to the burst release in the control cement without microparticles. However, the addition of 15% w/w microparticles to the cement decreased its ultimate compressive strength below the ISO 5833 standard minimum threshold of 70MPa [27]. In our study, we used a direct incubation method where we incubated our DOX-NP cement samples directly with the cells submersed in the cell medium, similar to Ozben *et al*. [50], which may be more indicative of the *in vivo* application.

In 3D culture, we assessed the metabolic activity and overall spheroid area in response to cement treatment. As tumors *in vivo* grow in a 3D environment, standard 2D cultures are unable to accurately simulate the native tumor microenvironment. Therefore, more physiologically relevant 3D models, such as spheroids, are increasingly being used for drug screening [51, 52]. Spheroids are characterized by proliferative (outer), quiescent (middle), and necrotic (inner) cell layers due to nutrient, gas, and pH gradients. They also generally possess higher drug resistances compared to standard monolayer cultures [53]. For the MDA-MB-231 cell line, we observed a similar trend in 3D as we did in 2D culture, where all the DOX-NP cement samples significantly reduced inhibited the metabolic activity of the spheroids. MDA-MB-231 spheroids have been shown to form loose spheroids and have similar drug sensitivities in 2D as they do in 3D culture [54]. Indeed, the MDA-MB-231 spheroids demonstrate an abundant proliferative zone as the tumor cells migrated outwards along the collagen matrix. The area of spheroids treated with the control blank and 0µM cements increased over the period of 7 days, whereas the DOX-NP cement treatments inhibited cell migration completely. A necrotic core is visible in the center of the control spheroids, as expected, due to the limited availability of nutrients and gas exchange in the core. With DOX-NP cement treatments, MDA-MB-231 spheroids showed dead cells and a reduced cell density as seen on phase contrast imaging. The C4-2B spheroids were more resistant to the DOX-NP cement treatment, compared to 2D culture. Mosaad *et al*. evaluated the metabolic activity of C4-2B spheroids in response to docetaxel and found that they were more resistant compared to 2D culture [55]. In contrast to the MDA-MB-231 spheroids, the C4-2B spheroids did not display an abundant outgrowth of cells. However, DOX-NP cement treatments reduced the size of the spheroids compared to control cement. Live/Dead imaging revealed a significant red fluorescent zone in the C4-2B spheroids treated with DOX-NP cement after 7 days. Spheroids treated with a free DOX concentration of 2µM in cell medium did not display the same degree of ethidium accumulation (red signal) in cells at the core. As the metabolic activity of the C4-2B spheroids was not drastically different between the treatment groups, it is unlikely that the fluorescent red signal is associated with dead cells from the Live/Dead staining. Rather, it may reflect the accumulation of DOX within the spheroid, which displays a red autofluorescence. An RFP filter cube was used for the fluorescence microscopy, which can also detect DOX [56]. Based on the DOX release kinetics, an estimated 1.5µM concentration should have been achieved in the cell medium by day 7. However, as the medium was not changed for the duration of the experiments, the pH drop may have increased the release kinetics leading to a higher dose of DOX and thereby causing more DOX accumulation in the spheroids. Furthermore, the metabolic activity of control C4-2B spheroids does not significantly increase over the course of the 7-day experiment, nor does the area increase as much as it does for the MDA-MB-231 spheroids. Future experiments could focus on determining the actual DOX uptake under these experimental conditions.

In comparing 2D and 3D cultures, a difference in the metabolic activities can be seen between the culture methods for the control spheroids. At day 7, MDA-MB-231 spheroids demonstrated a significantly higher metabolic activity in 3D culture, whereas C4-2B spheroids were significantly less active in 3D culture. Although there was no statistically significant difference between culture models for the DOX-NP cement treatment, the relative inhibition of MDA-MB-231 cells was higher in 3D culture, whereas it was higher in 2D culture for C4-2B cells. C4-2B spheroids have been reported to undergo moderate growth for the first 5-7 days, followed by a plateau in the proliferation rate [55]. Another prostate cancer cell line, PC3, has been shown to remain quiescent and proliferate slowly in 3D spheroid culture [57]. In contrast, spheroids formed with the highly invasive MDA-MB-231 cell line have been shown to grow rapidly over a period of 10 days [58]. As DOX targets rapidly dividing cells, it is expected that less proliferative or quiescent cells would be less affected by it [59]. This demonstrates the importance of 3D culture models for adequate high-throughput drug screening applications.

The limitations of the present study lie predominantly in the methods of predicting DOX release kinetics and the usage of cell lines. Predicting DOX release kinetics *in vivo* is challenging due to the differences in temperature, pH, medium used (e.g., PBS), etc. As release rates are determined by diffusion, the presence of concentration gradients can impact elution rates. In our assays, half the solution volume was replenished at each time point. In the *in vivo* case, however, DOX can diffuse into the entire organism, which may accelerate the release. Further experiments with various sink conditions are required to characterize release rates as a function of sink volumes and fractional replenishments. Established cell lines were used in this study for *in vitro* experiments, which may not accurately reflect spine metastatic cells. Future *in vitro* studies utilizing patient-derived spine metastatic cells in 3D culture would accurately predict the clinical success of our augmented PMMA cement. Nevertheless, our *in vitro* assessment in 2D and 3D culture showcases the efficacy of the DOX-NP cement for inhibiting cancer cell activity and migration. To our knowledge, no previous study has investigated the addition of chemotherapy-loaded mesoporous silica NPs to PMMA bone cement. Furthermore, we assess the *in vitro* efficacy of PMMA cement in 3D spheroid culture for the first time. Future work may assess combination chemotherapeutics for improved efficacy, various sizes of NPs and their functionalization for a better control of drug release profiles, as well as additional additives such as nanofibers to improve mechanical strength.

## Acknowledgments

AS received the CIHR CGS-M and FRQS doctoral scholarships. MEC received the CIHR postdoctoral fellowship award. DHR is a FRQS Junior 2 Research Scholar.

## References

[1] O. Barzilai, C. G. Fisher, and M. H. Bilsky, “State of the Art Treatment of Spinal Metastatic Disease,” Neurosurgery, vol. 82, no. 6, pp. 757–769, Jun 1 2018, doi: 10.1093/neuros/nyx567.

[2] R. Harel and L. Angelov, “Spine metastases: current treatments and future directions,” Eur J Cancer, vol. 46, no. 15, pp. 2696–707, Oct 2010, doi: 10.1016/j.ejca.2010.04.025.

[3] T. Liang, Y. Wan, X. Zou, X. Peng, and S. Liu, “Is surgery for spine metastasis reasonable in patients older than 60 years?,” Clin Orthop Relat Res, vol. 471, no. 2, pp. 628–39, Feb 2013, doi: 10.1007/s11999-012-2699-3.

[4] D. M. Sciubba and Z. L. Gokaslan, “Diagnosis and management of metastatic spine disease,” Surg Oncol, vol. 15, no. 3, pp. 141–51, Nov 2006, doi: 10.1016/j.suronc.2006.11.002.

[5] R. J. Rothrock et al., “Survival Trends After Surgery for Spinal Metastatic Tumors: 20-Year Cancer Center Experience,” Neurosurgery, vol. 88, no. 2, pp. 402–412, Jan 13 2021, doi: 10.1093/neuros/nyaa380.

[6] V. Kurisunkal, A. Gulia, and S. Gupta, “Principles of Management of Spine Metastasis,” Indian J Orthop, vol. 54, no. 2, pp. 181–193, Apr 2020, doi: 10.1007/s43465-019-00008-2.

[7] B. G. Bate, N. R. Khan, B. Y. Kimball, K. Gabrick, and J. Weaver, “Stereotactic radiosurgery for spinal metastases with or without separation surgery,” J Neurosurg Spine, vol. 22, no. 4, pp. 409–15, Apr 2015, doi: 10.3171/2014.10.SPINE14252.

[8] E. P. Howell et al., “Total en bloc resection of primary and metastatic spine tumors,” Ann Transl Med, vol. 7, no. 10, p. 226, May 2019, doi: 10.21037/atm.2019.01.25.

[9] K. Hayashi and H. Tsuchiya, “The role of surgery in the treatment of metastatic bone tumor,” Int J Clin Oncol, Feb 28 2022, doi: 10.1007/s10147-022-02144-6.

[10] J. H. Healey, F. Shannon, P. Boland, and G. R. DiResta, “PMMA to stabilize bone and deliver antineoplastic and antiresorptive agents,” Clin Orthop Relat Res, no. 415 Suppl, pp. S263–75, Oct 2003, doi: 10.1097/01.blo.0000093053.96273.ee.

[11] J. M. Cloyd, F. L. Acosta, Jr., M. Y. Polley, and C. P. Ames, “En bloc resection for primary and metastatic tumors of the spine: a systematic review of the literature,” Neurosurgery, vol. 67, no. 2, pp. 435–44; discussion 444-5, Aug 2010, doi: 10.1227/01.NEU.0000371987.85090.FF.

[12] B. Roedel et al., “Has the percutaneous vertebroplasty a role to prevent progression or local recurrence in spinal metastases of breast cancer?,” J Neuroradiol, vol. 42, no. 4, pp. 222–8, Jul 2015, doi: 10.1016/j.neurad.2014.02.004.

[13] N. Sundaresan, A. Rothman, K. Manhart, and K. Kelliher, “Surgery for solitary metastases of the spine: rationale and results of treatment,” Spine (Phila Pa 1976), vol. 27, no. 16, pp. 1802–6, Aug 15 2002, doi: 10.1097/00007632-200208150-00021.

[14] A. Ismat et al., “Antibiotic cement coating in orthopedic surgery: a systematic review of reported clinical techniques,” J Orthop Traumatol, vol. 22, no. 1, p. 56, Dec 23 2021, doi: 10.1186/s10195-021-00614-7.

[15] L. M. Mensah and B. J. Love, “A meta-analysis of bone cement mediated antibiotic release: Overkill, but a viable approach to eradicate osteomyelitis and other infections tied to open procedures,” Mater Sci Eng C Mater Biol Appl, vol. 123, p. 111999, Apr 2021, doi: 10.1016/j.msec.2021.111999.

[16] V. Wall, T. H. Nguyen, N. Nguyen, and P. A. Tran, “Controlling Antibiotic Release from Polymethylmethacrylate Bone Cement,” Biomedicines, vol. 9, no. 1, Jan 1 2021, doi: 10.3390/biomedicines9010026.

[17] A. Lilikakis and M. P. Sutcliffe, “The effect of vancomycin addition to the compression strength of antibiotic-loaded bone cements,” Int Orthop, vol. 33, no. 3, pp. 815–9, Jun 2009, doi: 10.1007/s00264-008-0521-3.

[18] P. Hernigou et al., “Methotrexate diffusion from acrylic cement. Local chemotherapy for bone tumours,” J Bone Joint Surg Br, vol. 71, no. 5, pp. 804–11, Nov 1989, doi: 10.1302/0301-620X.71B5.2584251.

[19] M. A. Rosa et al., “Acrylic cement added with antiblastics in the treatment of bone metastases. Ultrastructural and in vitro analysis,” J Bone Joint Surg Br, vol. 85, no. 5, pp. 712–6, Jul 2003.

[20] S. S. Phull, A. R. Yazdi, M. Ghert, and M. R. Towler, “Bone cement as a local chemotherapeutic drug delivery carrier in orthopedic oncology: A review,” J Bone Oncol, vol. 26, p. 100345, Feb 2021, doi: 10.1016/j.jbo.2020.100345.

[21] J. A. Handal, N. C. Tiedeken, G. E. Gershkovich, J. A. Kushner, B. Dratch, and S. P. Samuel, “Polyethylene glycol improves elution properties of polymethyl methacrylate bone cements,” J Surg Res, vol. 194, no. 1, pp. 161–6, Mar 2015, doi: 10.1016/j.jss.2014.10.051.

[22] G. A. Funk, J. C. Burkes, K. A. Cole, M. N. Rahaman, and T. E. McIff, “Antibiotic Elution and Mechanical Strength of PMMA Bone Cement Loaded With Borate Bioactive Glass,” J Bone Jt Infect, vol. 3, no. 4, pp. 187–196, 2018, doi: 10.7150/jbji.27348.

[23] J. A. Mendez, G. A. Abraham, M. del Mar Fernandez, B. Vazquez, and J. San Roman, “Self-curing acrylic formulations containing PMMA/PCL composites: properties and antibiotic release behavior,” J Biomed Mater Res, vol. 61, no. 1, pp. 66–74, Jul 2002, doi: 10.1002/jbm.10142.

[24] S. C. Shen, K. Letchmanan, P. S. Chow, and R. B. H. Tan, “Antibiotic elution and mechanical property of TiO2 nanotubes functionalized PMMA-based bone cements,” J Mech Behav Biomed Mater, vol. 91, pp. 91–98, Mar 2019, doi: 10.1016/j.jmbbm.2018.11.020.

[25] K. Letchmanan et al., “Mechanical properties and antibiotic release characteristics of poly(methyl methacrylate)-based bone cement formulated with mesoporous silica nanoparticles,” J Mech Behav Biomed Mater, vol. 72, pp. 163–170, Aug 2017, doi: 10.1016/j.jmbbm.2017.05.003.

[26] C. Kweon, A. C. McLaren, C. Leon, and R. McLemore, “Amphotericin B delivery from bone cement increases with porosity but strength decreases,” Clin Orthop Relat Res, vol. 469, no. 11, pp. 3002–7, Nov 2011, doi: 10.1007/s11999-011-1928-5.

[27] E. L. Cyphert, N. Kanagasegar, N. Zhang, G. D. Learn, and H. A. von Recum, “PMMA Bone Cement Composite Functions as an Adjuvant Chemotherapeutic Platform for Localized and Multi-Window Release during Bone Reconstruction,” Macromol Biosci, vol. 22, no. 5, p. e2100415, May 2022, doi: 10.1002/mabi.202100415.

[28] C. F. Thorn et al., “Doxorubicin pathways: pharmacodynamics and adverse effects,” Pharmacogenet Genomics, vol. 21, no. 7, pp. 440–6, Jul 2011, doi: 10.1097/FPC.0b013e32833ffb56.

[29] K. Krukiewicz and J. K. Zak, “Biomaterial-based regional chemotherapy: Local anticancer drug delivery to enhance chemotherapy and minimize its side-effects,” Mater Sci Eng C Mater Biol Appl, vol. 62, pp. 927–42, May 2016, doi: 10.1016/j.msec.2016.01.063.

[30] R. M. Sabio, A. B. Meneguin, T. C. Ribeiro, R. R. Silva, and M. Chorilli, “New insights towards mesoporous silica nanoparticles as a technological platform for chemotherapeutic drugs delivery,” Int J Pharm, vol. 564, pp. 379–409, Jun 10 2019, doi: 10.1016/j.ijpharm.2019.04.067.

[31] J. Slane, J. Vivanco, J. Meyer, H. L. Ploeg, and M. Squire, “Modification of acrylic bone cement with mesoporous silica nanoparticles: effects on mechanical, fatigue and absorption properties,” J Mech Behav Biomed Mater, vol. 29, pp. 451–61, Jan 2014, doi: 10.1016/j.jmbbm.2013.10.008.

[32] S. M. Khaled, P. A. Charpentier, and A. S. Rizkalla, “Physical and mechanical properties of PMMA bone cement reinforced with nano-sized titania fibers,” J Biomater Appl, vol. 25, no. 6, pp. 515–37, Feb 2011, doi: 10.1177/0885328209356944.

[33] S. C. Shen, W. K. Ng, Z. Shi, L. Chia, K. G. Neoh, and R. B. Tan, “Mesoporous silica nanoparticle-functionalized poly(methyl methacrylate)-based bone cement for effective antibiotics delivery,” J Mater Sci Mater Med, vol. 22, no. 10, pp. 2283–92, Oct 2011, doi: 10.1007/s10856-011-4397-1.

[34] S. C. Shen, W. K. Ng, Y. C. Dong, J. Ng, and R. B. Tan, “Nanostructured material formulated acrylic bone cements with enhanced drug release,” Mater Sci Eng C Mater Biol Appl, vol. 58, pp. 233–41, Jan 1 2016, doi: 10.1016/j.msec.2015.08.011.

[35] J. Shen, Q. He, Y. Gao, J. Shi, and Y. Li, “Mesoporous silica nanoparticles loading doxorubicin reverse multidrug resistance: performance and mechanism,” Nanoscale, vol. 3, no. 10, pp. 4314–22, Oct 5 2011, doi: 10.1039/c1nr10580a.

[36] E. L. Cyphert, G. D. Learn, D. W. Marques, C. Y. Lu, and H. A. von Recum, “Antibiotic Refilling, Antimicrobial Activity, and Mechanical Strength of PMMA Bone Cement Composites Critically Depend on the Processing Technique,” ACS Biomater Sci Eng, vol. 6, no. 7, pp. 4024–4035, Jul 13 2020, doi: 10.1021/acsbiomaterials.0c00305.

[37] H. Li et al., “Cisplatin and doxorubicin dual-loaded mesoporous silica nanoparticles for controlled drug delivery,” RCS Advances, vol. 6, no. 96, pp. 94160–94169, 2016.

[38] L. Feng, Z. Dong, D. Tao, Y. Zhang, and Z. Liu, “The acidic tumor microenvironment: a target for smart cancer nano-theranostics,” National Science Review, vol. 5, no. 2, pp. 269–286, 2018.

[39] V. Estrella et al., “Acidity generated by the tumor microenvironment drives local invasion,” Cancer Res, vol. 73, no. 5, pp. 1524–35, Mar 1 2013, doi: 10.1158/0008-5472.CAN-12-2796.

[40] A. Alkaraki et al., “Enhancing chemosensitivity of wild-type and drug-resistant MDA-MB-231 triple-negative breast cancer cell line to doxorubicin by silencing of STAT 3, Notch-1, and beta-catenin genes,” Breast Cancer, vol. 27, no. 5, pp. 989–998, Sep 2020, doi: 10.1007/s12282-020-01098-9.

[41] J. Ni et al., “Epithelial cell adhesion molecule (EpCAM) is associated with prostate cancer metastasis and chemo/radioresistance via the PI3K/Akt/mTOR signaling pathway,” Int J Biochem Cell Biol, vol. 45, no. 12, pp. 2736–48, Dec 2013, doi: 10.1016/j.biocel.2013.09.008.

[42] J. Han et al., “Chemoresistance in the Human Triple-Negative Breast Cancer Cell Line MDA-MB-231 Induced by Doxorubicin Gradient Is Associated with Epigenetic Alterations in Histone Deacetylase,” J Oncol, vol. 2019, p. 1345026, 2019, doi: 10.1155/2019/1345026.

[43] M. Tonak et al., “HDAC inhibitor-loaded bone cement for advanced local treatment of osteosarcoma and chondrosarcoma,” Anticancer Res, vol. 34, no. 11, pp. 6459–66, Nov 2014.

[44] K. Koto, H. Murata, Y. Sawai, E. Ashihara, M. Horii, and T. Kubo, “Cytotoxic effects of zoledronic acid-loaded hydroxyapatite and bone cement in malignant tumors,” Oncol Lett, vol. 14, no. 2, pp. 1648–1656, Aug 2017, doi: 10.3892/ol.2017.6355.

[45] C. C. Chiang, M. K. Hsieh, C. Y. Wang, W. H. Tuan, and P. L. Lai, “Cytotoxicity and cell response of preosteoblast in calcium sulfate-augmented PMMA bone cement,” Biomed Mater, vol. 16, no. 5, Aug 19 2021, doi: 10.1088/1748-605X/ac1ab5.

[46] J. Fang, J. Shen, W. Jiang, W. Dong, and Z. Hu, “Cytotoxicity of polymethyl methacrylate cement on primary cultured metastatic spinal cells,” Molecular & Cellular Toxicology, vol. 12, no. 2, pp. 125–132, 2016.

[47] M. F. Moreau, D. Chappard, M. Lesourd, J. P. Montheard, and M. F. Basle, “Free radicals and side products released during methylmethacrylate polymerization are cytotoxic for osteoblastic cells,” J Biomed Mater Res, vol. 40, no. 1, pp. 124–31, Apr 1998, doi: 10.1002/(sici)1097-4636(199804)40:1<124::aid-jbm14>3.0.co;2-o.

[48] F. Greco, L. de Palma, N. Specchia, S. Jacobelli, and C. Gaggini, “Polymethylmethacrylate-antiblastic drug compounds: an in vitro study assessing the cytotoxic effect in cancer cell lines--a new method for local chemotherapy of bone metastasis,” Orthopedics, vol. 15, no. 2, pp. 189–94, Feb 1992, doi: 10.3928/0147-7447-19920201-13.

[49] G. Maccauro et al., “Methotrexate-added acrylic cement: biological and physical properties,” J Mater Sci Mater Med, vol. 18, no. 5, pp. 839–44, May 2007, doi: 10.1007/s10856-006-0036-7.

[50] H. Ozben, L. Eralp, G. Baysal, A. Cort, N. Sarkalkan, and T. Ozben, “Cisplatin loaded PMMA: mechanical properties, surface analysis and effects on Saos-2 cell culture,” Acta Orthop Traumatol Turc, vol. 47, no. 3, pp. 184–92, 2013, doi: 10.3944/aott.2013.2828.

[51] A. A. Fitzgerald, E. Li, and L. M. Weiner, “3D Culture Systems for Exploring Cancer Immunology,” Cancers (Basel), vol. 13, no. 1, Dec 28 2020, doi: 10.3390/cancers13010056.

[52] M. Zanoni et al., “3D tumor spheroid models for in vitro therapeutic screening: a systematic approach to enhance the biological relevance of data obtained,” Sci Rep, vol. 6, p. 19103, Jan 11 2016, doi: 10.1038/srep19103.

[53] M. P. Carvalho, E. C. Costa, S. P. Miguel, and I. J. Correia, “Tumor spheroid assembly on hyaluronic acid-based structures: A review,” Carbohydr Polym, vol. 150, pp. 139–48, Oct 5 2016, doi: 10.1016/j.carbpol.2016.05.005.

[54] Y. Imamura et al., “Comparison of 2D- and 3D-culture models as drug-testing platforms in breast cancer,” Oncol Rep, vol. 33, no. 4, pp. 1837–43, Apr 2015, doi: 10.3892/or.2015.3767.

[55] E. O. Mosaad, K. F. Chambers, K. Futrega, J. A. Clements, and M. R. Doran, “The Microwell-mesh: A high-throughput 3D prostate cancer spheroid and drug-testing platform,” Sci Rep, vol. 8, no. 1, p. 253, Jan 10 2018, doi: 10.1038/s41598-017-18050-1.

[56] T. Yildiz, R. Gu, S. Zauscher, and T. Betancourt, “Doxorubicin-loaded protease-activated near-infrared fluorescent polymeric nanoparticles for imaging and therapy of cancer,” Int J Nanomedicine, vol. 13, pp. 6961–6986, 2018, doi: 10.2147/IJN.S174068.

[57] A. Y. Hsiao et al., “Microfluidic system for formation of PC-3 prostate cancer co-culture spheroids,” Biomaterials, vol. 30, no. 16, pp. 3020–7, Jun 2009, doi: 10.1016/j.biomaterials.2009.02.047.

[58] D. P. Saraiva, A. T. Matias, S. Braga, A. Jacinto, and M. G. Cabral, “Establishment of a 3D Co-culture With MDA-MB-231 Breast Cancer Cell Line and Patient-Derived Immune Cells for Application in the Development of Immunotherapies,” Front Oncol, vol. 10, p. 1543, 2020, doi: 10.3389/fonc.2020.01543.

[59] S. Y. van der Zanden, X. Qiao, and J. Neefjes, “New insights into the activities and toxicities of the old anticancer drug doxorubicin,” FEBS J, vol. 288, no. 21, pp. 6095–6111, Nov 2021, doi: 10.1111/febs.15583.

